# Genomic prediction of stalk lodging resistance and the associated intermediate phenotypes in maize using whole-genome resequence and multi-environmental data

**DOI:** 10.1101/2025.04.06.647466

**Authors:** Caique Machado e Silva, Bharath Kunduru, Norbert Bokros, Kaitlin Tabaracci, Yusuf Oduntan, Manwinder S. Brar, Rohit Kumar, Christopher J. Stubbs, Maicon Nardino, Christopher S. McMahan, Seth DeBolt, Daniel J. Robertson, Rajandeep S. Sekhon, Gota Morota

## Abstract

Breeding for stalk lodging resistance is of paramount importance to maintain and improve maize yield and quality and meet increasing food demand. The integration of environmental, phenotypic, and genotypic information offers the opportunity to develop genomic prediction strategies that can improve the genetic gain for complex traits such as stalk lodging. However, implementation of genomic predictions for stalk lodging resistance has been sparse primarily due to the lack of reliable and reproducible phenotyping strategies. In this study, we measured 10 traits related to stalk lodging resistance obtained from a novel phenotyping platform on approximately 31,000 individual stalks. These traits were combined with environmental information and whole-genome resequence data to investigate the predictive ability of different single and multi-environment genomic prediction models. In total, 555 maize inbred lines from the Wisconsin diversity panel were evaluated in four environments. The multi-environment models more than doubled the prediction accuracy compared to the single-environment model for most traits, particularly when predicting lines in a sparse testing design. Predictive correlations for stalk bending strength and stalk flexural stiffness, a non-destructive method for assessment of stalk lodging resistance, were moderately high and ranged between 0.32 to 0.89 and 0.26 to 0.88, respectively. In contrast, rind thickness was the most difficult trait to predict. Our results show that the use of multi-environmental data could improve genomic prediction accuracy for stalk lodging resistance and its intermediate phenotypes. This study will serve as a first step toward genetic improvement and the development of maize varieties resistant to stalk lodging.

**Core Ideas:**

- The DARLING platform was successfully used to collect lodging resistance-related phenotypes
- Proportion of variation explained by genotype by environment interaction was not negligible
- Genomic predictions for lodging resistance phenotypes ranged from moderate to high
- Accounting for genotype by environment interaction was found to be important for improved predictive performance

**Plain language summary:** Stalk lodging - when maize stalks break or fall over before harvest - can seriously reduce crop yields. Breeding maize that resists lodging is important to ensure reliable food production. We tested about 31,000 individual stalks for 10 traits related to lodging resistance using a new phenotyping system. These traits were combined with environmental information and whole-genome resequence data to investigate the predictive ability of single and multi-environment genomic prediction models. By analyzing 555 maize lines, we found that using data from multiple environments improved genomic prediction accuracy by more than twice as much as using data from a single environment. Traits such as bending strength and flexural stiffness were easier to predict, while rind thickness was more difficult. These results show that combining genetic, environmental, and phenotypic data can help breeders more accurately select maize plants with stronger stalks, leading to better, more resilient crops.

## Introduction

Maize yield and quality have increased tremendously worldwide (Andorf et al., 2019), mainly due to the elucidation of the basis of heterosis and the development of hybrid technology (Shull, 1908, 1909, 1910; East, 1936). However, continued increases in yield to meet future global food demand (Falcon et al., 2022) are hampered by several challenges that negatively affect maize production (Anderson et al., 2020). Lodging, the permanent displacement of a plant from its vertical position, causes 2-43% yield losses worldwide (Kunduru et al., 2023). Lodging is a complex trait strongly influenced by genetic, physiological, morphological, and environmental factors (Ye et al., 2016; Erndwein et al., 2020; Liu et al., 2020; Xue et al., 2020; Robertson et al., 2022; Bokros et al., 2024). Lodging occurs due to either the inability of the roots to keep plants erect and implanted in the soil (root lodging) or mechanical failure of the stalk (stalk lodging) (Sekhon et al., 2020). Both root and stalk lodging make crops difficult to harvest and significantly reduce grain yield and quality, resulting in billions of dollars in losses worldwide each year (Kunduru et al., 2023). As the world climate continues to deteriorate, losses due to lodging are expected to increase and seriously threaten food security (Rezaei et al., 2023). Understanding the genetic architecture of lodging resistance and applying this knowledge in breeding programs is a key part of preventing such losses in the future.

Reliable phenotyping of crop germplasm for stalk lodging resistance is necessary for genetic improvement and variety development. However, assessment of stalk lodging resistance under field conditions has remained a major challenge for breeding and genetic studies due to lack of reliable and reproducible phenotyping strategies (Kunduru et al., 2023; Stubbs et al., 2023). Traditionally, stalk lodging in maize has been assessed under field conditions by counting the number of naturally lodged plants. However, this phenotyping method is not reproducible as the natural stalk lodging patterns are highly influenced by environment and, under optimal conditions, the genotypes possessing poor stalk lodging qualities may not present the phenotype. Furthermore, the data produced by this approach are limited to a binary output (i.e., lodged or not lodged) which is not ideal for genetic analysis of a quantitative trait. Alternatively, many indirect measures of stalk lodging resistance, hereafter referred to as *intermediate phenotypes*, are often used as proxies for estimating stalk lodging resistance (Zuber and Grogan, 1961; Thompson, 1963). Recently, a novel approach to quantitatively measure stalk lodging resistance was proposed by employing the DARLING (Device for Assessing Resistance to Lodging IN Grains) (Cook et al., 2019; Seegmiller et al., 2020; Stubbs et al., 2020a; DeKold and Robertson, 2023). The DARLING is a phenotyping platform capable of reproducing stalk failure patterns similar to those observed under natural conditions (Robertson et al., 2015a) and thus provides both a reliable and reproducible strategy to assess stalk lodging in maize (Robertson et al., 2015b; Sekhon et al., 2020; Stubbs et al., 2020c). Importantly, besides providing stalk bending strength, a destructive measurement, the DARLING can also be used to produce a non-destructive measure of lodging resistance in the form of stalk flexural stiffness (Cook et al., 2019; Erndwein et al., 2020). The latter is particularly important for artificial selection of ideal genotypes in breeding programs as the desirable plants can be used for genetic crosses and harvesting seed.

Availability of the DARLING platform offers opportunities for genetic and breeding studies aimed to improve stalk lodging resistance in maize. Phenotypic estimates of stalk lodging resistance produced by the DARLING and other novel and traditional intermediate phenotypes can be combined with whole-genome resequence data to predict the performance of genotypes in field trials. In particular, genomic prediction estimates the effect of markers or variants across the genome to predict the genomic estimated breeding values of selection candidates that have not yet been phenotyped (Bernardo, 1994; Meuwissen et al., 2001). When phenotypes are collected in multiple environments, the predictive performance of genomic prediction models can be enhanced by accounting for the interaction between genotypes and environments (i.e., G × E) (Burgueño et al., 2012; Jarquín et al., 2014; Lopez-Cruz et al., 2015; Cuevas et al., 2017; Dalsente Krause et al., 2020). An advantage of genomic prediction is that it does not require prior knowledge of the exact location of the causal variants or genes.

Genomic prediction studies of stalk lodging resistance and the associated intermediate phenotypes in maize are scarce and limited to single trait and single environment studies with a small recombinant inbred line population (Liu et al., 2020). In the current study, we integrated multiple novel and traditional intermediate phenotypes related to stalk lodging resistance collected on a large set of dent maize inbred lines evaluated in multiple environments with high density whole-genome resequence data to perform genomic prediction analysis. The main objective of this study was to compare the predictive ability of different single- and multi-environment genomic prediction models for maize stalk lodging resistance-related traits taking into account the G × E interaction. The findings of this study lay a foundation for the implementation of a genomic selection paradigm aimed at improving stalk lodging resistance within maize.

## Materials and Methods

### Genetic materials and field experiments

A subset of the maize Wisconsin diversity panel (Mazaheri et al., 2019) consisting of 555 inbred maize lines was evaluated at two locations during the 2020 and 2021 summer seasons in the USA (Table 1). In Pendleton, SC, experiments were conducted at the Simpson Research and Education Center of Clemson University. In Lexington, KY, experiments were conducted at the Spindletop Farm of the University of Kentucky. The combinations of locations and years were used to define a total of four environments. Experiments were designed as a randomized complete block design with two replications. Plots consisted of single row of 6.1 m with 0.76 m length spacing. The target population density was 70,000 plants ha^*−*1^. We followed standard agronomic practices as described elsewhere (Kunduru et al., 2023).

**Table 1:** Geographical description of the locations where the experiments were conducted during the 2020 and 2021 summer seasons.

| Location | Year | Environment | Latitude | Longitude | Average Altitude |
| --- | --- | --- | --- | --- | --- |
| Pendleton, SC, USA | 2020 | C20 | 34° 39' N | 82° 47' W | 260 m |
| Pendleton, SC, USA | 2021 | C21 | 34° 39' N | 82° 47' W | 260 m |
| Lexington, KY, USA | 2020 | K20 | 38° 2' N | 84° 30' W | 276 m |
| Lexington, KY, USA | 2021 | K21 | 38° 2' N | 84° 30' W | 276 m |

### Phenotypic data

Phenotypes evaluated in this study are considered suitable indicators of stalk lodging resistance in the field (Sekhon et al., 2020; Kunduru et al., 2023; Tabaracci et al., 2024). At physiological maturity, approximately 38-42 days after anthesis, representative non-lodged plants from each plot were assessed (Kunduru et al., 2023) to obtain data on intermediate phenotypes at plant and internode levels (Table 2). These include bending strength, flexural stiffness, ear height, major diameter, minor diameter, plant height, and rind thickness. Phenotypes assessed across multiple internodes include, major diameter, minor diameter, and rind thickness, were recorded on the bottom internode (BI, corresponding to the first elongated internode above the ground level) and the ear internode (EI, corresponding to the internode immediately below the primary-ear bearing node). A total of ten intermediate phenotypes were analyzed in this study.

**Table 2:** Description of the traits analyzed to evaluate stalk lodging resistance of maize inbred lines in four environments.

| Trait | Category | Sampling unit | Unit |
| --- | --- | --- | --- |
| Bending strength | Structural | Stalk | N mm<sup>2</sup> |
| Flexural stiffness | Structural | Stalk | N mm<sup>2</sup> |
| Ear height | Morphological | Stalk | cm |
| Major diameter | Geometric | Internode | mm |
| Minor diameter | Geometric | Internode | mm |
| Plant height | Morphological | Stalk | cm |
| Rind thickness | Geometric | Internode | mm |

Two reliable indicators of stalk lodging resistance in maize, bending strength and flexural stiffness, were measured using the DARLING (Cook et al., 2019). Briefly, the DARLING consists of a footplate and a vertical arm with an attached force sensor. During testing, the device is aligned with the designated stalk at a selected height and a point load is applied to the stalk. The applied force and deflection angle of the stalk is recorded (Stubbs et al., 2023). These data are then used to calculate bending strength and flexural stiffness, where bending strength is the maximum bending moment the stalk can tolerate before breaking (Robertson et al., 2015b; Stubbs et al., 2020b; Sekhon et al., 2020) and flexural stiffness is a measurement of the resistance of the stalk to deflection (Stubbs et al., 2023; Kunduru et al., 2023).

Major diameter, minor diameter, and rind thickness are considered as geometric traits. Both the BI and EI of the genotypes were subjected to rind penetration tests to determine rind thickness (Stubbs et al., 2020a; Cook et al., 2020; Seegmiller et al., 2020). Rind penetration tests were performed using an Instron universal testing machine by forcing a probe through the internode at a rate of 25 mm s^*−*1^. The resulting force-displacement curve was then analyzed using a custom MATLAB algorithm to calculate rind thickness and minor diameter. The major diameters of the internodes were measured with a caliper. Finally, plant height and ear height, which are considered morphological traits, were collected under field conditions as described in our previous study (Kunduru et al., 2023).

The phenotypic data were unbalanced because not all lines were evaluated across the four environments and because of exclusion of some data due to strict quality control measures followed during phenotyping and data processing. Quality control measures were also performed to facilitate the construction of the cross-validation designs described below. Inbred lines with missing records in at least two environments were excluded from the analysis. Phenotypic values of inbred lines with missing records in one environment were imputed using the adjusted means of the 20 closely related lines according to the genomic relationship matrix (VanRaden, 2008). The number of inbred lines analyzed for each trait is shown in Table 3.

**Table 3:** Number of inbred lines and number of single nucleotide polymorphisms (SNPs) used in the study after quality control. BI: bottom internode and EI: ear internode

| Trait | Number of lines | Number of SNPs |
| --- | --- | --- |
| Bending strength | 326 | 11,582,742 |
| Ear height | 325 | 11,569,562 |
| Flexural stiffness | 327 | 11,600,634 |
| Major diameter-BI | 326 | 11,571,925 |
| Major diameter-EI | 326 | 11,571,925 |
| Minor diameter-BI | 326 | 11,594,337 |
| Minor diameter-EI | 326 | 11,600,634 |
| Plant height | 325 | 11,569,562 |
| Rind thickness-BI | 314 | 11,497,391 |
| Rind thickness-EI | 325 | 11,582,361 |

### Environmental data

Environmental information for the experiment conducted in Lexington was obtained from the Blue Grass Airport Station, Lexington (https://www.wunderground.com/). Environmental information for the experiment conducted in Pendleton was obtained from the National Climatic Data Center database (https://www.ncei.noaa.gov/cdo-web/). A summary of the environmental data is presented in Table 4. These environmental data were used in multi-environment genomic prediction models described below.

**Table 4:** Summary of the environmental information retrieved for four environments during conduction of experiments. C20: Clemson University 2020 data, K20: University of Kentucky 2020 data, C21: Clemson University 2021 data, and K21: University of Kentucky 2021 data.

| Environmental information | Unit | C20 | K20 | C21 | K21 |
| --- | --- | --- | --- | --- | --- |
| Maximum temperature | $^{\circ}$ C | 35.55 | 33.33 | 33.89 | 33.33 |
| Minimum temperature | $^{\circ}$ C | -1.11 | -3.89 | -3.89 | -1.11 |
| Total precipitation | mm | 906.78 | 547.12 | 760.98 | 667.77 |

### Genomic data

Whole-genome resequencing data for all the inbred lines included in the present study was obtained from the public domain (Grzybowski et al., 2023). Briefly, whole-genome resequencing data from 1,276 previously published maize samples (Chia et al., 2012; Unterseer et al., 2014; Brandenburg et al., 2017; Wang et al., 2017; Bukowski et al., 2018; Kistler et al., 2018; Wang et al., 2020; Qiu et al., 2021; Chen et al., 2022) were combined with resequencing data from 239 inbred maize lines to generate a high-density genome-wide marker set with more than 46 million high-confidence single nucleotide polymorphism (SNP) variants. The original resequenced data contained 14,793,812 SNP markers. A quality control process removed markers with correlation ≥ 0.99, minor allele frequency ≤ 5%, and missing genotype rate ≥ 10%. Since the number of inbred lines for each phenotype varied slightly due to the phenotypic quality control described above, the number of SNPs also varied. A minimum of 11,000,000 SNPs were used for each phenotype. The number of SNPs used for each phenotype is shown in Table 3.

### Phenotypic analyses

All statistical analyses described below were performed in R version 4.4.2. (R Core Team, 2024). Phenotypic data were analyzed to derive adjusted means using the SpATS function in the SpATS R package (Rodriguez-Alvarez et al., 2019). Genomic prediction models were implemented using the BGLR R package (Pérez and de Los Campos, 2014). Inference was based on 20,000 Markov Chain Monte Carlo iterations with a burn-in of 8,000 and a thinning rate of five.

#### Best linear unbiased estimation

Best linear unbiased estimators (BLUE) of genotypes were estimated by analyzing the phenotypic data. Field plot coordinates were used to correct for spatial heterogeneity using a P-spline model (Rodriguez-Alvarez et al., 2018), which uses the tensor product P-splines to model large- and small-scale spatial trends:

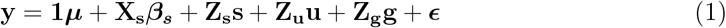

where **y** is the vector of phenotypic observations; **1** is a vector of ones; ***µ*** is the overall mean; **X**_**s**_***β***_***s***_ is the fixed non-penalized term and **Z**_**s**_**s** is the random penalized term [**s** ∼ *N* (0, **S**)] of the smooth spatial surface mixed model, i.e., *f* (*r, c*) = **X**_**s**_***β***_***s***_+**Z**_**s**_**s**. *f* (*r, c*) can be decomposed into a sum of linear components and unviariate and bivariate smooth functions as follows :

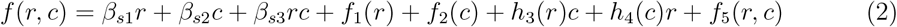

where *β*_*s*1_*r* is the linear trend across rows; *β*_*s*2_*c* is the linear trend across columns; *β*_*s*3_*rc* is the linear interaction trend between rows and columns; *f*_1_(*r*) is the smooth trend across rows; *f*_2_(*c*) is the smooth trend across columns; *h*_3_(*r*)*c* and *h*_4_(*c*)*r* are the two linear-by-smooth interaction terms, where the slope of a linear trend along one covariate (c or r) is allowed to vary smoothly as function of the other covariate; and *f*_5_(*r, c*) is the smooth-by-smooth interaction between column and row trends. Also, the spatial covariance matrix **S** is the direct sum of the matrices 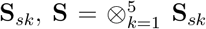, where each parameter of *sk* depends on a specific smoothing parameter (*λ*_*sk*_), which can be estimated via restricted maximum likelihood as the ratio between the residual variance and the corresponding variance of spatial effects, i.e., 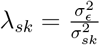. Finally, **u** is the vector that comprises the mutually independent sub-vectors of row and column effects accounting for discontinuous field variation, **g** is the vector of genotype fixed effects; **Z**_**u**_ and **Z**_**g**_ are the corresponding incidence matrices; and 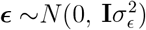 is a vector of residuals. The BLUE of the genotypes was used in all subsequent single- and multi-environment genomic prediction models.

### Single-environment genomic prediction

A single-environment genomic best linear unbiased prediction (GBLUP) model was fitted to derive the genomic estimated breeding values of the inbred lines. The GBLUP model can be represented as follows:

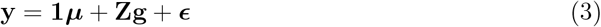

where **y** is a vector of BLUE; 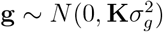 is a vector of additive genetic effects; 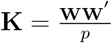 is the genomic relationship matrix of the inbred lines (VanRaden, 2008); **W** is the centered and standardized SNP matrix; *p* is the total number of SNPs; and 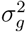 is the additive genetic variance.

### Multi-environment genomic prediction

Multi-environment models used in this work are extensions of the single-environment GBLUP and capture information across correlated environments. In the first fitted model, genetic and environmental effects are described as linear functions of SNPs and environmental covariates (ECs), respectively (Jarquín et al., 2014). This model is hereafter referred to as the genotype by covariate interaction model (G × C). In G × C, the G × E interaction is accounted for by modeling the interaction between the genomic similarities (i.e., genomic relationship matrix) and the environmental similarities derived from ECs. The second model fitted was the marker × environment interaction model (M × E) (Lopez-Cruz et al., 2015). This model decomposes the genetic variance into components that are stable across environments (main effects) and environment-specific components (interaction effects). The last model fitted was the G × E interaction kernel model (Cuevas et al., 2017), which can be interpreted as a multi-trait model (MTM) that considers each environment as a different trait. The MTM model considers an effect that simultaneously estimates the genetic and G × E interaction effects (MTM1). This model has also been extended to include an additional component (MTM2) that is designed to capture genetic effects that are not fully captured by MTM1. Detailed descriptions of the fitted multi-environment models are given below.

### Genotype by covariate interaction

The fitted G × C interaction model is (Jarquín et al., 2014)

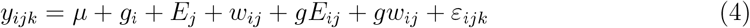

where *y*_*ijk*_ is the phenotypic observation; *µ* is the overall mean; 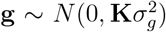 is additive genetic effects of the *i*th line; 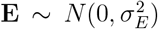 is the environmental effect of the *j*th environment (*j* = 1, *···*, 4); 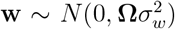 is the EC effect; **Ω** describes similarities between environments listed in Table 4 and computed similar to the genomic relationship matrix **K**, but using ECs instead of gene contents;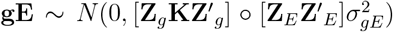 is the interaction between the genetic and environmental effects; **Z**_*g*_ is the incidence matrix for the genetic effect; **Z**_*E*_ is the incidence matrix for the environmental effect; ∘ is the Hadamard product; 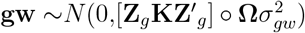 is the interaction between genetic and EC effects; and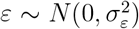 is the residual. Also, 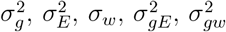, and 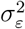 are corresponding variance components.

### Marker × environment interaction

The M × E interaction model explicitly regresses a vector of phenotypes on the SNP marker matrix as follows (Lopez-Cruz et al., 2015):

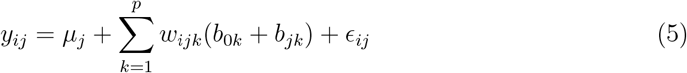

where *y*_*ij*_ is the phenotype of the *i*th genotype in the *j*th environment; *µ*_*j*_ is the overall mean of the *j*th environment; *w*_*ijk*_ is the centered and standardized *k*th marker covariate; 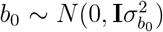 is the common effect of markers across environments; 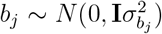 is the specific effect of markers unique to each environment; and 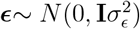 is the residual. In matrix notation, equation (5) can be expressed as

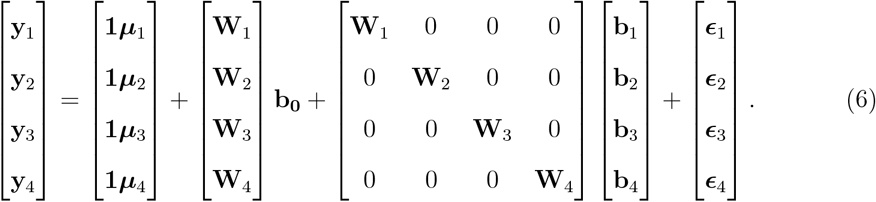

where subscripts 1 *···*4 denote the four environments.

The model described in equation (5) is equivalent to the M × E ridge regression BLUP. Alternatively, one can fit the M × E interaction model by assuming covariance structures for the genetic effect as follows (M × E interaction GBLUP).

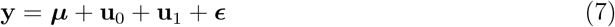

where *µ* is the overall mean; 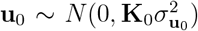 is the common main environmental effect; **u**_1_ ∼ *N* (0, **K**_1_) is the environment-specific effect; and 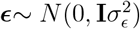 is the residual. Here

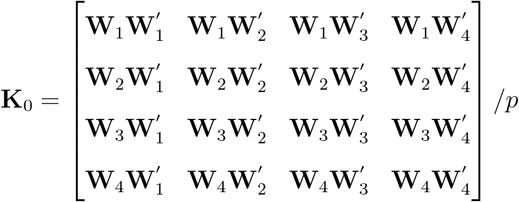

and

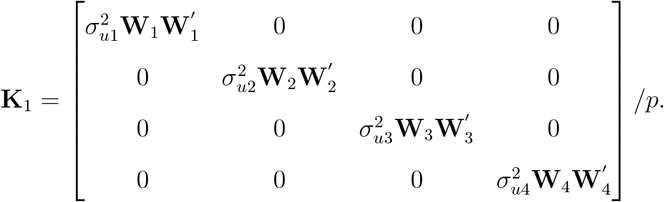

In the current work, the GBLUP parameterization of the M × E model was used.

### Multi-trait model

In MTM, the genetic effect **u** is estimated considering simultaneously the similarity between genotypes and the variance-covariance matrix between environments. MTM is expressed as follows (MTM1) (Cuevas et al., 2017):

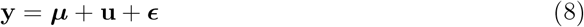

where **u** ∼ *N* (0, **G**) is the genetic effect; 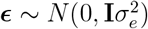 is the residual; and **G** is the Kronecker product between the unstructured genetic variance-covariance matrix of environments (**E**) and the genomic relationship matrix (**K**).

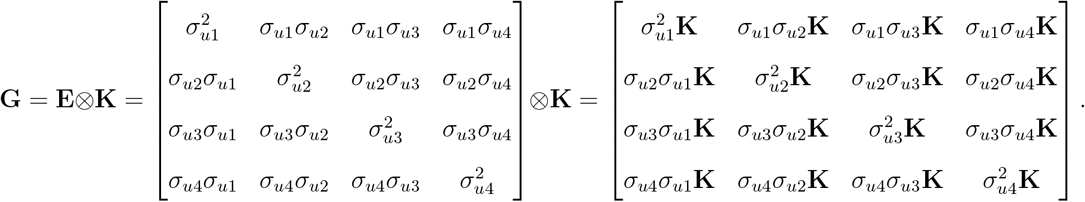

In many cases, the component **u** may not capture all existing genetic variability due to imperfect linkage disequilibrium between markers and quantitative trait loci or model misspecification (Cuevas et al., 2017). In such a case, it is reasonable to extend MTM1 by considering an additional component **f**, which is expected to capture genetic variability not explained by **u**. Such a model can be expressed as (MTM2):

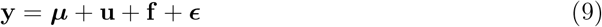

where **f** ∼ *N* (0, **Q**) is the genetic effect that captures the genetic variability not accounted for by **u** and **Q** is the Kronecker product between the unstructured variance-covariance matrix of **F** and an identity matrix **I**.

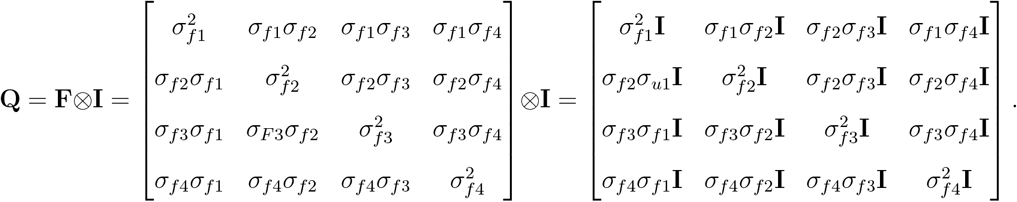

### Cross-validation schemes

The predictive ability of the single-environment GBLUP model was evaluated using a repeated random subsampling cross-validation scheme with 100 replications. The data set was divided into 80% training and 20% testing. The predictive ability was measured as the Pearson’s correlation between the predicted and observed values in the testing set. For the multi-environment models, the predictive ability was evaluated considering two different cross-validation scenarios (CV1 and CV2). CV1 mimics the case where the breeder is interested in predicting the performance of lines that have not been observed in any environment, i.e., newly developed lines. CV2 mimics the situation where the breeder is interested in predicting the performance of lines that have been observed in some environments but are missing in others, i.e., sparse testing design. In both scenarios, the data set was randomly partitioned into 80% training and 20% testing. The predictive ability of the models was also assessed as the Pearson’s correlation between the predicted and observed values in the testing set.

## Results

### Genomic heritability estimates

Moderate to low genomic heritability estimates were obtained from the single-environment GBLUP analysis, with values ranging from 0.22 (rind thickness-EI, K20) to 0.54 (minor diameter-BI, C21) (Figure 1). Genomic heritability estimates for a given trait varied somewhat among environments. For example, the estimate for rind thickness-BI in C20 was 0.47, while the estimates for the same trait in C21 and K20 were 0.27. This observation suggests that, besides the underlying genetics, the environmental conditions play an important role in the genetic expression of the intermediate phenotypes associated with stalk lodging resistance. Therefore, integration of information across environments is of paramount importance for genetic evaluation and prediction of stalk lodging resistance in maize. Genomic heritability estimates for traits derived from the DARLING, bending strength and flexural stiffness, were moderate and higher compared to some other traits (e.g.,plant height, rind thickness-BI, and rind thickness-EI), especially in C20 (0.41 for bending strength and 0.47 for flexural stiffness) and C21 (0.46 for bending strength and 0.42 for flexural stiffness). This evidence suggests that bending strength and flexural stiffness could be used routinely to evaluate germplasm for stalk lodging resistance in maize breeding programs.

**Figure 1:**
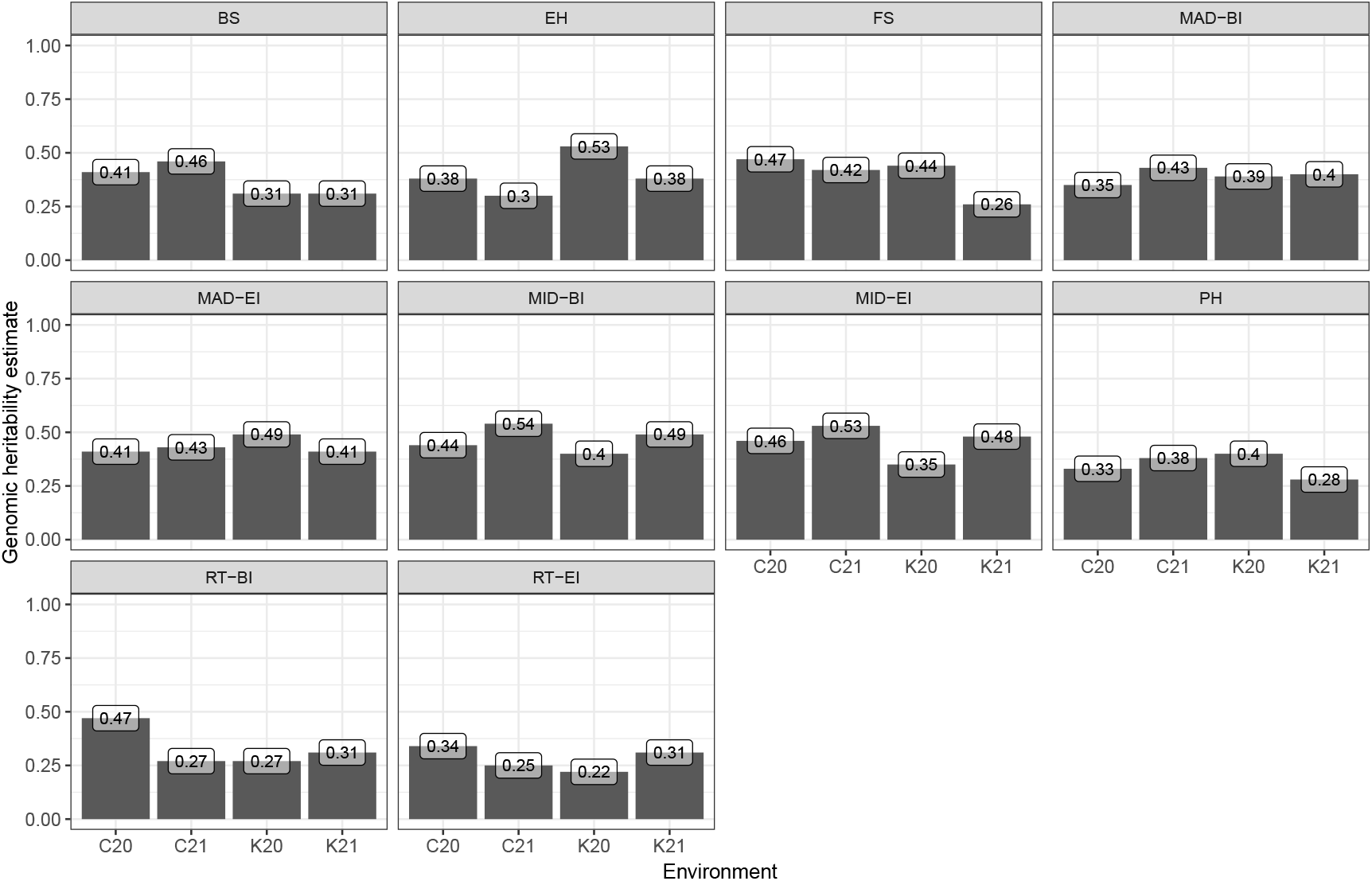
Genomic heritability estimates of stalk lodging resistance-related traits evaluated in Clemson University 2020 (C20), Clemson University 2021 (C21), University of Kentucky 2020 (K20), and University of Kentucky 2021 (K21). BS: Bending strength; EH: Ear height; FS: Flexural stiffness; MAD: Major diameter; MID: Minor diameter; PH: Plant height; RT: Rind thickness; BI: bottom internode; and EI: ear internode.

### Genetic correlation estimates

Because multi-environment models leverage information from correlated environments, a greater extent of genetic correlation between environments is generally desired. In this study, positive genetic correlations between the four environments using MTM were observed for all traits (Figure 2). In addition, genetic correlation estimates were high in most cases (≥ 0.7), except for rind thickness-BI and rind thickness-EI. For these traits, genetic correlation estimates between environments were low to moderate, and had the lowest genomic heritability estimates, as shown previously (Figure 1).

**Figure 2:**
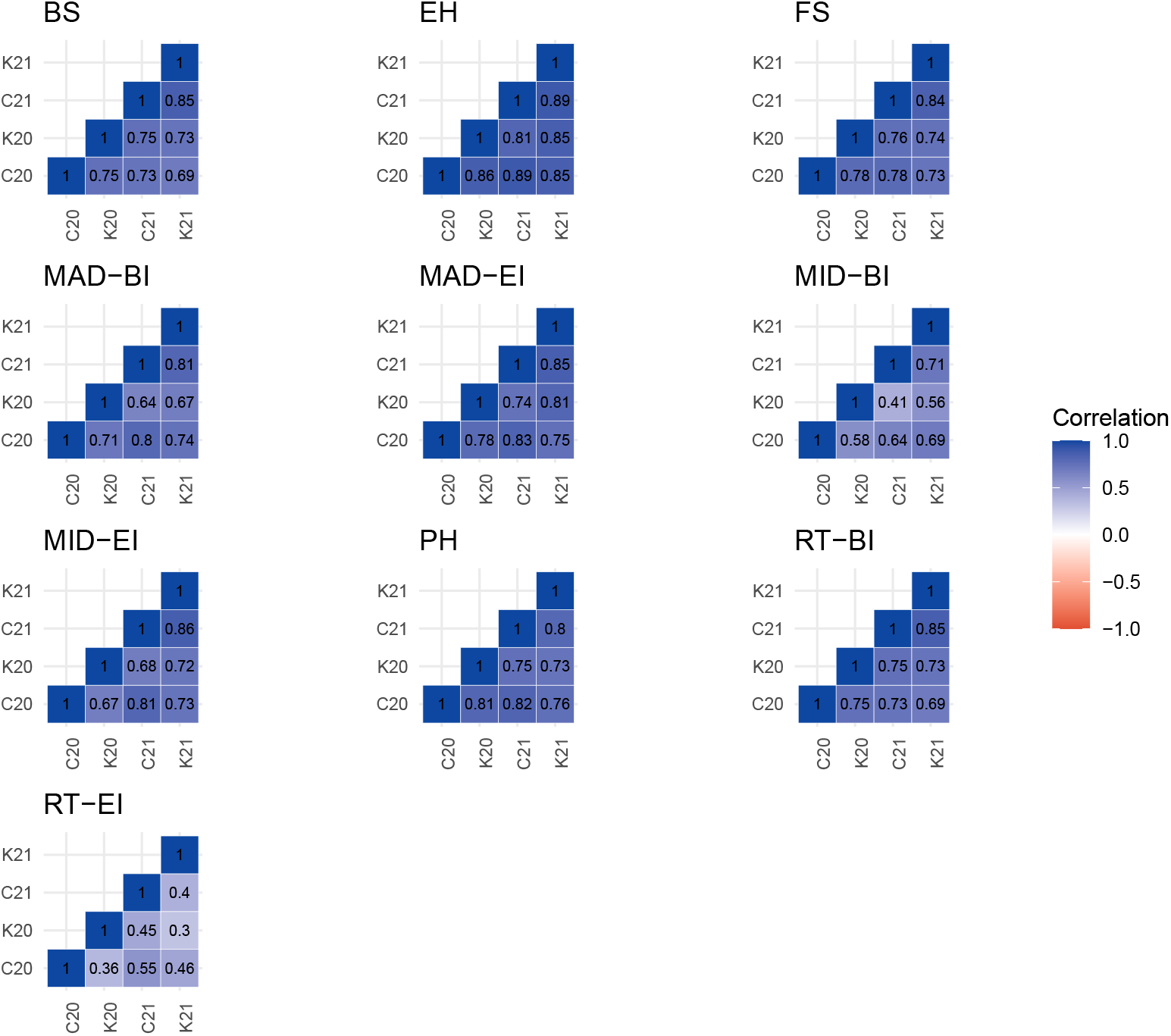
Genetic correlation estimates between environments for stalk lodging resistance-related traits. BS: Bending strength; EH: Ear height; FS: Flexural stiffness; MAD: Major diameter; MID: Minor diameter; PH: Plant height; RT: Rind thickness; BI: bottom internode; and EI: ear internode.

### Multi-environment model parameters

Proportions of the variance components for the environmental, environmental covariate, genetic, and interaction effects estimated from the G × C interaction model are shown in Figure 3. For most traits, the genetic effect contributed a sizable proportion of the total variance. Since selection acts on the heritable portion of phenotypic variation, large estimates of genetic variance components are desirable. However, for some traits, the residual variance accounted for a large proportion of the phenotypic variation (e.g., rind thickness-BI and rind thickness-EI). This is consistent with the low to moderate single-environment genomic heritability estimates obtained for these traits. Estimates of environmental variance contributed a smaller fraction of the total variance observed compared to genetic variance, except for rind thickness-BI. In addition, the proportion of variance accounted for by environmental covariates was small. Thus, it is possible that the environmental covariates used in the study did not capture all the variation between environments and that additional components may need to be included in the model. Estimates of the interaction terms (G × E and G × EC) were small but not negligible, suggesting that it is important to include them in the prediction models.

**Figure 3:**
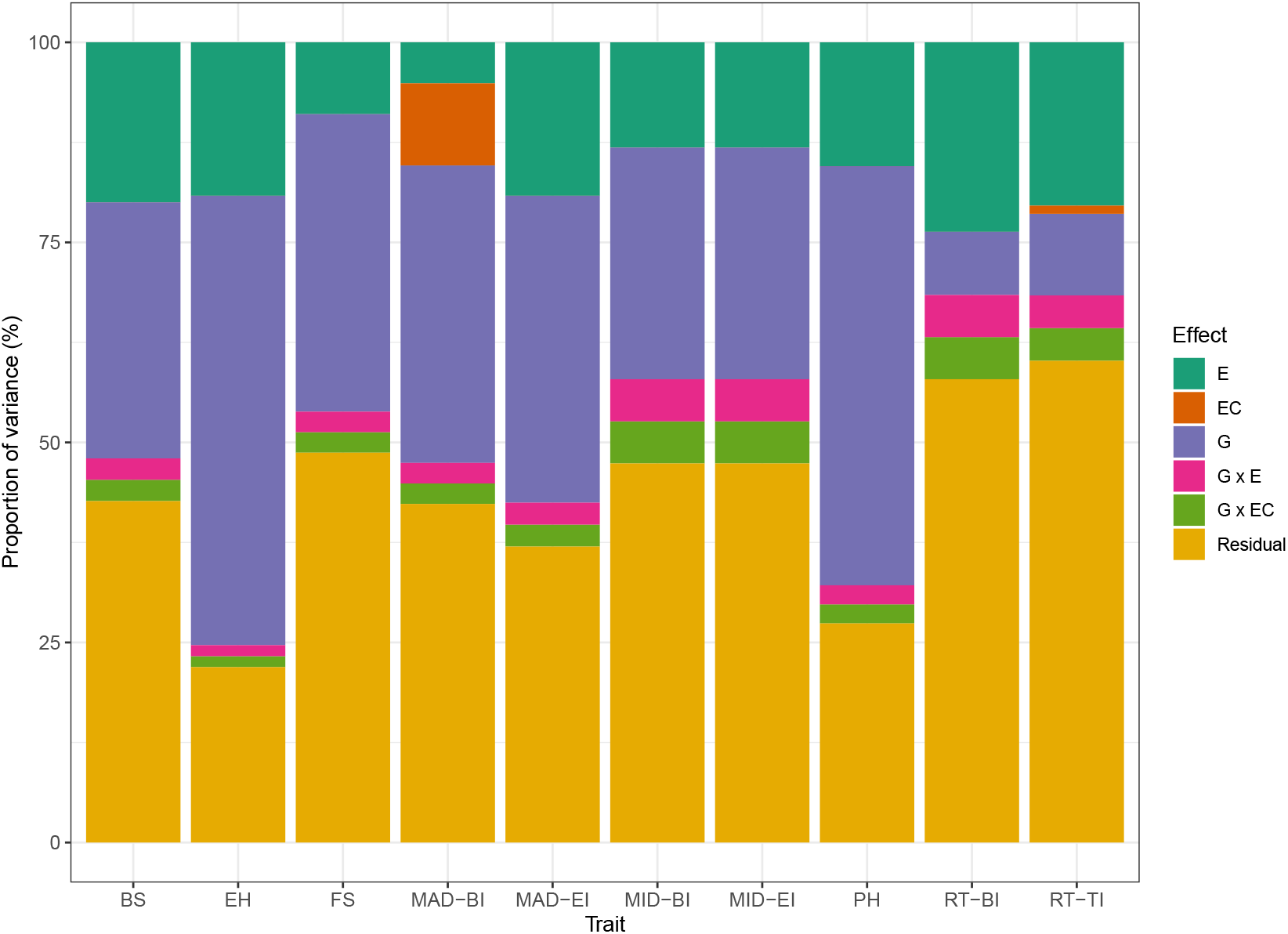
Proportion of environmental (E), environmental covariate (EC), genetic (G), G x E, and G x EC interaction variances estimated from the G × C interaction model for a set of stalk lodging resistance-related traits. BS: Bending strength; EH: Ear height; FS: Flexural stiffness; MAD: Major diameter; MID: Minor diameter; PH: Plant height; RT: Rind thickness; BI: bottom internode; and EI: ear internode.

Proportions of the variance components due to the main genetic, environment-specific, and residual effects estimated from the M × E interaction model are shown in Figure 4. The main genetic effect accounted for a sizable proportion of the total variance observed for most traits. It is worth noting that the variance component of the main genetic effect was higher for the traits that had a high correlation between environments (Figure 2). In contrast, traits that had a low estimate of genetic correlation between environments (e.g., rind thickness-BI and rind thickness-EI) had a greater contribution of the environment-specific effect represented by the interaction term. The interaction term in the M × E model played a larger role in the total variation observed for most traits than in the G × C model. In addition, for all traits, the residual variances had a smaller contribution to the total variance compared to those estimated from the G × C model.

**Figure 4:**
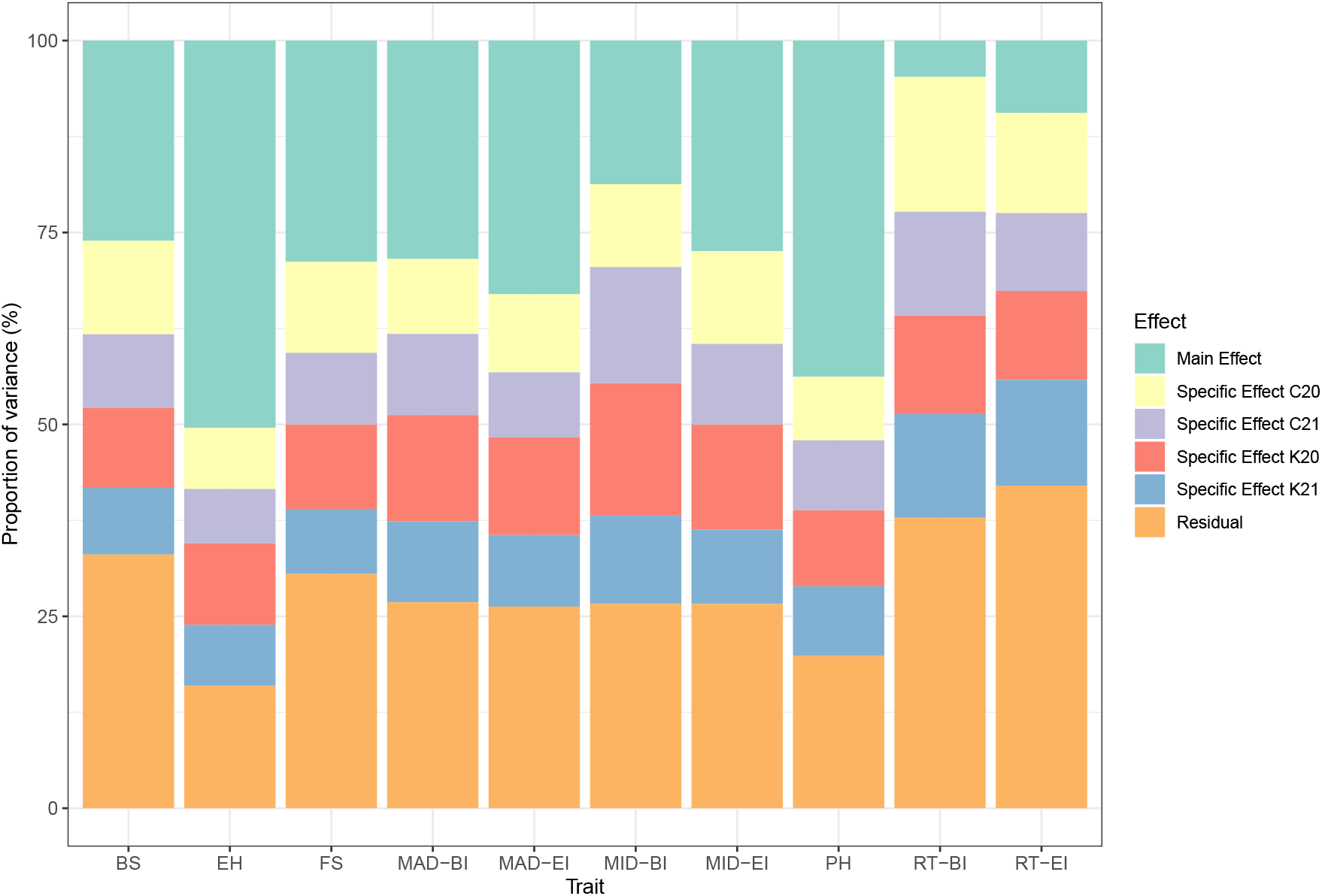
Proportion of main genetic, specific and residual variances obtained from the M x E model for a set of stalk lodging resistance-related traits. BS: Bending strength; EH: Ear height; FS: Flexural stiffness; MAD: Major diameter; MID: Minor diameter; PH: Plant height; RT: Rind thickness; BI: bottom internode; and EI: ear internode.

### Predictive ability

Predictive ability of the single-environment GBLUP and the multi-environment models evaluated using the two cross-validation schemes are shown in Figure 5. The predictive ability of the single-environment GBLUP ranged from 0.12 (plant height, C20) to 0.42 (bending strength, C21). No significant improvement was observed when multi-environment models were fitted to predict newly developed lines (CV1). In this scenario, the predictive ability ranged from 0.10 (plant height, C20) to 0.44 (bending strength, C21) in G × C; 0.11 (plant height, C20) to 0.42 (BS, C21) in M × E; 0.10 (plant height, C20) to 0.42 (bending strength, C21) in MTM1; and 0.11 (plant height, C20) to 0.44 (bending strength, C21) in MTM2.

**Figure 5:**
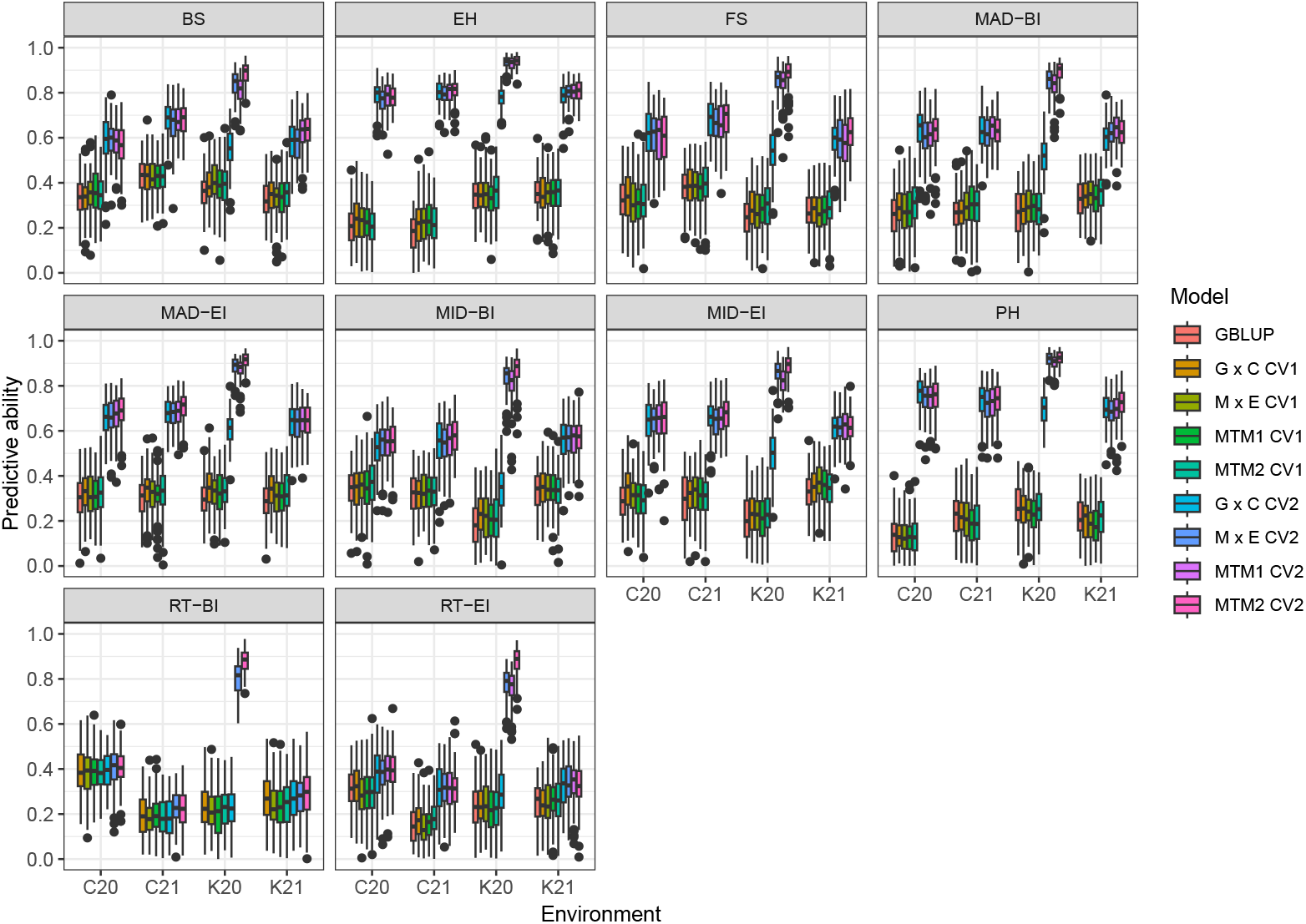
Predictive ability of the single-trait single-environment genomic best linear unbiased prediction (GBLUP) model and various multi-environment genomic prediction models (G × C, M × E, MTM1, and MTM2) using the two cross-validation schemes (CV1 and CV2) for stalk lodging resistance-related traits evaluated in Clemson University 2020 (C20), Clemson University 2021 (C21), University of Kentucky 2020 (K20), and University of Kentucky 2021 (K21). BS: Bending strength; EH: Ear height; FS: Flexural stiffness; MAD: Major diameter; MID: Minor diameter; PH: Plant height; RT: Rind thickness; BI: bottom internode; and EI: ear internode.

When the multi-environment models were fitted to predict the performance of lines in a sparse testing design (CV2), a notable increase in predictive ability was observed for most of the traits. In CV2, the predictive ability ranged from 0.20 (rind thickness-BI, C21) to 0.80 (ear height, C21) in G × C; 0.22 (rind thickness-BI, C21) to 0.93 (ear height, K20) in M × E; 0.21 (rind thickness-BI, C21) to 0.93 (ear height, K20) in MTM1; and 0.21 (rind thickness-BI, C21) to 0.94 (ear height, K20) in MTM2. In general, the multi-environment models had similar predictive ability across the environments except for the G × C interaction model in K20, where the predictive correlations were smaller compared to the other multi-environment models. Notably, even in CV2, no big improvement was observed for rind thickness-BI and rind thickness-EI and some multi-environment models were only able to increase the predictive ability in K20 for these traits. On average, the multi-environment models in CV2 improved predictive ability by 107% (G × C), 135% (M × E), 132% (MTM1), and 139% (MTM2) over the single-environment GBLUP model.

## Discussion

The development of superior genetic material in plant breeding programs relies on the accurate phenotypic evaluation of genotypes (inbred lines, hybrids, segregating populations, etc.) in diverse environments. Novel quantitative trait phenotyping for stalk lodging resistance has emerged as a promising tool for the evaluation of various traits in economically important crops and with several advantages over the existing methods. In particular, evaluation of the intermediate phenotypes associated with stalk lodging resistance in maize lines using novel albeit standardized phenotyping pipelines offers the opportunity to characterize a wide range of genotypes in a rapid and accurate manner (Cook et al., 2019; Tabaracci et al., 2024). Moreover, reductions in genotyping costs and advances in next-generation sequencing techniques have allowed us to obtain a large number of variants for many individuals in a cost-effective manner (Elshire et al., 2011; Poland and Rife, 2012). Combining high-dimensional phenotypic and high density genotypic information in advanced quantitative genetic frameworks (Morota et al., 2022) can provide insight into the genetic architecture of stalk lodging resistance and predict the performance of unobserved genotypes for rapid genetic enhancement.

Breeding for complex traits, such as stalk lodging resistance, is not an easy task. Because genes may be expressed differently across environments, statistical models that accommodate the complexity underlying the interaction between genotypes and environmental factors should be preferred. In general, GBLUP extended to multi-environment models borrows information from correlated environments. In this study, environments had moderate to high positive correlations with each other for most traits thus allowing us to improve the predictive ability in CV2 for most traits. The positive genetic correlations between environments can be explained by the relative temporal and geographic proximity of the four environments. Increasing the number of environments (seasons, years, locations, etc.) may potentially lead to a decrease in correlation estimates. In such a case, identifying non-complex G × E interaction patterns and grouping more correlated environments may be a better strategy for building experimental networks and targeting cultivar recommendations (Costa-Neto et al., 2020; Evangelista et al., 2021; Casagrande et al., 2024; Araújo et al., 2024).

The variance component estimates obtained from both the G × C and M × E interaction models show that the interaction terms are not negligible. For example, in the G × C model, the interaction terms (G × E and G × EC) accounted for up to 7.89% (rind thickness-BI) of the total observed variation. In the M × E interaction model, the interaction term (environment-specific effect) explained up to 57.43% (rind thickness-BI) of the total variation observed. Differences between these two models in the variance component estimation of the interaction terms were expected because G × C and M × E model the G × E interaction in different manners. In particular, G × C includes ECs capable of explaining variation across environments in addition to the random environmental effect. However, depending on the ECs used, they may not be able to successfully explain large proportions of such variation (Jarquín et al., 2014). In such a case, both the EC and G × EC variance component estimates will be low. In contrast, the variance components of the interaction terms in the M × E model are directly related to the correlation estimates, such that higher correlation estimates lead to larger contributions of the main genetic effects while lower estimates lead to larger contributions of the interaction terms as observed for rind thickness-BI and rind thickness-EI. This happens because the M × E interaction model imposes constraints on covariance structures such that covariances are forced to be positive and constant across environments (Lopez-Cruz et al., 2015).

Regardless of the model fitted, no notable improvements in predictive ability were observed when predicting newly developed lines (CV1). This is because predicting lines that have never been observed does not allow information to be borrowed across correlated environments. Instead, borrowing of information occurs only between related lines within environments. Conversely, when predictions were made considering a sparse testing design (CV2), the predictive ability increased for most traits. This observation is explained by the fact that prediction of a line in an environment allows borrowing of both within-environment and between-environment information as reported in previous studies (Burgueño et al., 2012; Peiffer et al., 2014; Jarquín et al., 2017; Dalsente Krause et al., 2020; Sabag et al., 2023). For rind thickness-BI and rind thickness-EI, however, the predictive ability did not increase significantly even under CV2 in most environments which may be caused by the low estimates of genomic heritability and genetic correlation. In such a scenario, an alternative potential strategy to increase the accuracy of these traits is to implement nonlinear Gaussian kernel models (Morota and Gianola, 2014; Bandeira e Sousa et al., 2017). Such models are able to capture small, complex interactions between markers and are expected to increase the predictive ability on the presence of epistasis.

The multi-environment genomic prediction models presented in this study have peculiarities that should be considered by breeders. The G × C interaction model avoids the problem of explicitly modeling all possible contrasts between markers and ECs by introducing a multiplicative model. Including interactions between markers and ECs improves predictive ability by increasing the proportion of variance accounted for by the model. However, ECs may not be able to explain all of the variation between environments and this fact may prevent a significant increase in predictive ability. This finding explains the slightly lower predictive ability estimates of G × C compared to the other models in general. In this case, the development and deployment of advanced environmental sensors in the field will help obtain environmental covariates that may be able to explain a greater proportion of phenotypes across environments. Finally, one of the limitations of G × C is that interactions can occur in many ways in practice, and this model works as an approximation to the unobserved real complex G × E interaction phenomena (Jarquín et al., 2014).

The M × E interaction model allows the genetic effect to be decomposed into effects that are common across environments and environment-specific effects (Lopez-Cruz et al., 2015; Sabag et al., 2023). This model may be of particular interest if breeders wish to select for overall performance and stability. However, due to the restrictions imposed on the covariance structures, the increase in predictive ability using the M × E model depends on the correlation between environments. As a consequence, environments that are highly correlated allow for higher predictive ability estimates, while poorly correlated environments lead to lower predictive abilities. This limitation may explain the slightly lower average predictive ability of M × E compared to MTM2.

To circumvent some of the drawbacks of the previous models, MTM simultaneously models genetic and environmental effects and is considered to be parsimonious (Cuevas et al., 2017). This model considers an unstructured environmental variance-covariance matrix and is expected to perform well even in environments with high G × E interaction and low correlation. In addition, MTM has the potential to be extended to account for genetic variation that may not be fully explained by marker effects. Our study showed that MTM2 had slightly better predictive performance than the other models that only considered genetic effects explained by markers. Therefore, MTM2 can be considered as the best fitting model in the current study and could be used to predict stalk lodging resistant genotypes in different environments. However, MTM is not without limitations and fitting such a model can be computationally challenging when the number of environments increases exponentially (Cuevas et al., 2017).

Genomic prediction of stalk lodging resistance in maize has often been overlooked, mainly due to the lack of reliable quantitative measures of lodging. A previous study performed genomic prediction of stalk lodging resistance in maize by considering the rind penetration resistance of the third stalk internode above the ground (Liu et al., 2020). One primary advantage of using bending strength or flexural stiffness to evaluate maize stalk lodging resistance is that they provide a holistic estimate of stalk lodging resistance at the plant-level as compared to other phenotypic measurements which only represents a single internode (e.g., rind thickness, rind penetration, diameter etc.). The morphological, geometric, and material properties of internodes vary across the stalk and, therefore, data collected on a single internode may not accurately represent the true lodging-related characteristics of the entire stalk (Kunduru et al., 2023; Oduntan et al., 2024). Internode-level phenotypes have other associated challenges as well. For example, rind thickness is notoriously difficult to quantify as it can change both along the length and circumference of a single internode and requires use of somewhat arbitrary thresholding techniques to determine (Oduntan et al., 2022). Whereas rind penetration does not account for key geometric properties of the stalk. In fact, it appears that selecting directly for rind penetration puts negative selection pressure on important geometric factors such as the moment of inertia that are key determinants of lodging resistance (Masole, 1993; Martin et al., 2004; Robertson et al., 2017; Stubbs et al., 2022; Oduntan et al., 2024). In summary, bending strength and flexural stiffness have been shown to be superior predictors of stalk lodging resistance and have higher heritability estimates compared to other internode-level phenotypes, including rind penetration resistance, and therefore, are ideal intermediate phenotypes to evaluate stalk lodging resistance (Robertson et al., 2016; Sekhon et al., 2020; Kunduru et al., 2023). While certain phenotyping strategies rely on cutting, transporting, and evaluating stalks in the laboratory, potentially adding data artifacts, both bending strength and flexural stiffness are collected in the field and, therefore, provide a more natural estimate of stalk lodging resistance at reduced cost. Furthermore, flexural stiffness is a non-destructive phenotype that allows the plant to be used simultaneously for phenotyping and genetic crosses, increasing the efficiency of breeding efforts.

The previous study performing genomic prediction of stalk lodging in maize (Liu et al., 2020) used recombinant inbred lines with a limited number of markers, whereas the current study used an unstructured population of whole-genome resequence data. It is important to note the long time required to establish a population for linkage mapping studies (e.g., recombinant inbred lines). In addition, the genetic basis of such a population is likely to be narrow, and for this reason the genomic prediction accuracy tends to be higher than that of an unstructured population because the lines are closely related. Besides bending strength and flexural stiffness, our study used a wide range of novel and traditional intermediate phenotypes and present a road map for using such phenotype in genomic selection endeavours. Therefore, depending upon resources available to the breeding program, one or more of these intermediate phenotypes can be used to implement genomic selection for stalk lodging resistance. Lastly, this study aimed to predict the performance of *per se* inbred lines. Further studies should focus on the prediction of single-cross hybrids (Dias et al., 2020; Barreto et al., 2024). In this case, in addition to the additive genetic effect, dominance effects should also be considered, as their inclusion has been shown to improve accuracy (Dias et al., 2018).

## Conclusion

Stalk lodging resistance in maize is a complex trait that is ultimately determined by numerous quantitative phenotypes that are challenging to acquire in a high throughput manner. To overcome these challenges, this study employed a novel phenotyping strategy and thereby demonstrated that genomic prediction can be successfully implemented to improve stalk lodging resistance in maize. Genomic selection models that use information from multiple environments are preferred over the single-trait single-environment model for predicting stalk lodging resistance. Predictive correlations for stalk bending strength and stalk flexural stiffness were moderately high and ranged between 0.32 to 0.89 and 0.26 to 0.88, respectively. Predicting phenotypes of newly developed inbred lines is more challenging than predicting the phenotypes of existing inbred lines in a sparse testing design. Finally, the integration of environmental data and high-dimensional genomic information offers the opportunity to improve genetic gains by increasing the accuracy of selection.

## Abbreviations

BI: bottom internode
BLUE: best linear unbiased estimates
CV: cross-validation
EI: ear internode
EC: environmental covariate
GBLUP: genomic best linear unbiased estimates
G × C: genotype by covariate interaction
MTM: multi-trait model
M × E: marker by environment interaction
SNP: single nucleotide polymorphism

## Author contributions

CMS: Conceptualization; Formal analysis, Methodology; Visualization; Writing-original draft. BK: Investigation; Data curation; Writing-review & editing. NB: Investigation; Data curation; Writing-review & editing. KT: Investigation; Data curation; Writing-review & editing. YO: Investigation; Data curation; Writing-review & editing. MSB: Investigation; Writing-review & editing. RK: Investigation; Writing-review & editing. CJS: Investigation; Data curation; Writing-review & editing. MN: Conceptualization; Writing-review & editing. CSM: Funding acquisition; Project administration; Writing-review & editing. SD: Funding acquisition; Project administration; Writing-review & editing. DJR: Funding acquisition; Project administration; Investigation, Data curation; Writing-review & editing. RSS: Funding acquisition; Project administration; Investigation; Data curation; Writing-review & editing. GM: Conceptualization; Methodology; Funding acquisition; Project administration; Supervision; Writing-original draft; Writing-review & editing.

## Acknowledgments

CMS acknowledge the Brazilian Federal Foundation for Support and Evaluation of Graduate Education (CAPES) for providing a scholarship to conduct research at Virginia Tech.

## Conflict of interest

The authors declare no conflicts of interest.

## Funding

This work was funded by the National Science Foundation EPSCoR grant #1826715 and in part by Virginia Tech.

## Notes

### Competing Interest Statement

The authors have declared no competing interest.

